# MscS inactivation and recovery are slow voltage-dependent processes sensitive to interactions with lipids

**DOI:** 10.1101/2023.05.08.539870

**Authors:** Madolyn Britt, Elissa Moller, Joseph Maramba, Andriy Anishkin, Sergei Sukharev

## Abstract

Mechanosensitive channel MscS, the major bacterial osmolyte release valve, shows a characteristic adaptive behavior. With a sharp onset of activating tension, the channel population readily opens, but under prolonged action of moderate near-threshold tension, it inactivates. The inactivated state is non-conductive and tension-insensitive, which suggests that the gate gets uncoupled from the lipid-facing domains. The kinetic rates for tension-driven opening-closing transitions are 4-6 orders of magnitude higher than the rates for inactivation and recovery. Here we show that inactivation is augmented and recovery is slowed down by depolarization. Hyperpolarization, conversely, impedes inactivation and speeds up recovery. We then address the question of whether protein-lipid interactions may set the rates and influence voltage dependence of inactivation and recovery. Mutations of conserved arginines 46 and 74 anchoring the lipid-facing helices to the inner membrane leaflet to tryptophans do not change the closing transitions, but instead change the kinetics of both inactivation and recovery and essentially eliminate their voltage-dependence. Uncharged polar substitutions (S or Q) for these anchors produce functional channels but increase the inactivation and reduce the recovery rates. The data suggest that it is not the activation and closing transitions, but rather the inactivation and recovery pathways that involve substantial rearrangements of the protein-lipid boundary associated with the separation of the lipid-facing helices from the gate. The discovery that hyperpolarization robustly assists MscS recovery indicates that membrane potential can regulate osmolyte release valves by putting them either on the ‘ready’ or ‘standby’ mode depending on the cell’s metabolic state.

## Introduction

MscS, the mechanosensitive channel of small conductance, is the main low-threshold osmolyte release valve regulating turgor pressure in most prokaryotes (Cox et al., 2018; Kung et al., 2010). Its molecular design was ‘successful’ in evolution, and multiple homologs of MscS are found in all clades of walled organisms, including higher plants (Hamilton et al., 2015). In *E. coli* and other bacteria, MscS usually acts in tandem with the high-threshold large-conductance mechanosensitive channel MscL (Levina et al., 1999; Naismith and Booth, 2012). Both channels gate directly in response to lateral tension in the lipid bilayer (Sukharev, 2002; Sukharev et al., 1993). Unlike MscL, however, MscS possesses the distinctive ability to inactivate, which renders it insensitive to mechanical stimuli and non-conductive. Inactivation is an important functional trait that helps to reduce the active population at low tensions and thus carefully dose membrane permeability and metabolic losses under mild hypotonic shocks (refs). Inactivation also plays a critical role in the termination of the massive permeability response to extreme shocks when both channels are engaged (Moller, 2023).

It is important to note that both the opening and inactivation transitions in MscS are driven by membrane tension, and both originate from the resting state. The ranges of preferred tensions are overlapping, and the maximum of inactivation probability takes place at tensions near the threshold of activation (Akitake et al., 2005; Cetiner et al., 2018). Previous studies have given estimations for the rates of the opening-closing transitions as well as the rates of inactivation-recovery. The former exhibit stronger tension dependence and higher transition rates. Patch-clamp recordings show that the rate of WT MscS activation varies steeply with tension, from estimated 10^−6^ s^-1^ at low (zero) tension to 10^5^ s^-1^ at ∼10 mN/m (Cetiner et al., 2018). Inactivation and recovery, in contrast, are notoriously slow, ranging between 0.07 and 0.15 s^-1^ (Cetiner et al., 2018; Kamaraju and Sukharev, 2008) for inactivation and between 0.5 and 1 s^-1^ for recovery (Belyy et al., 2010b; Kamaraju and Sukharev, 2008; Rowe et al., 2014). The spatiotemporal parameters estimated for MscS are presented in the gating scheme (Figure 1A). The two kinetically disparate conformational pathways follow different effective spatial scales: the in-plane expansion associated with opening (C→O) is estimated as 12-18 nm^2^ and the protein area change between the closed and inactivated states (C→I) is 8-9 nm^2^ (Akitake et al., 2005; Boer et al., 2011; Cetiner et al., 2018; Kamaraju et al., 2011) clearly indicating that the closed conformation must be the most compact.

**Figure 1.**
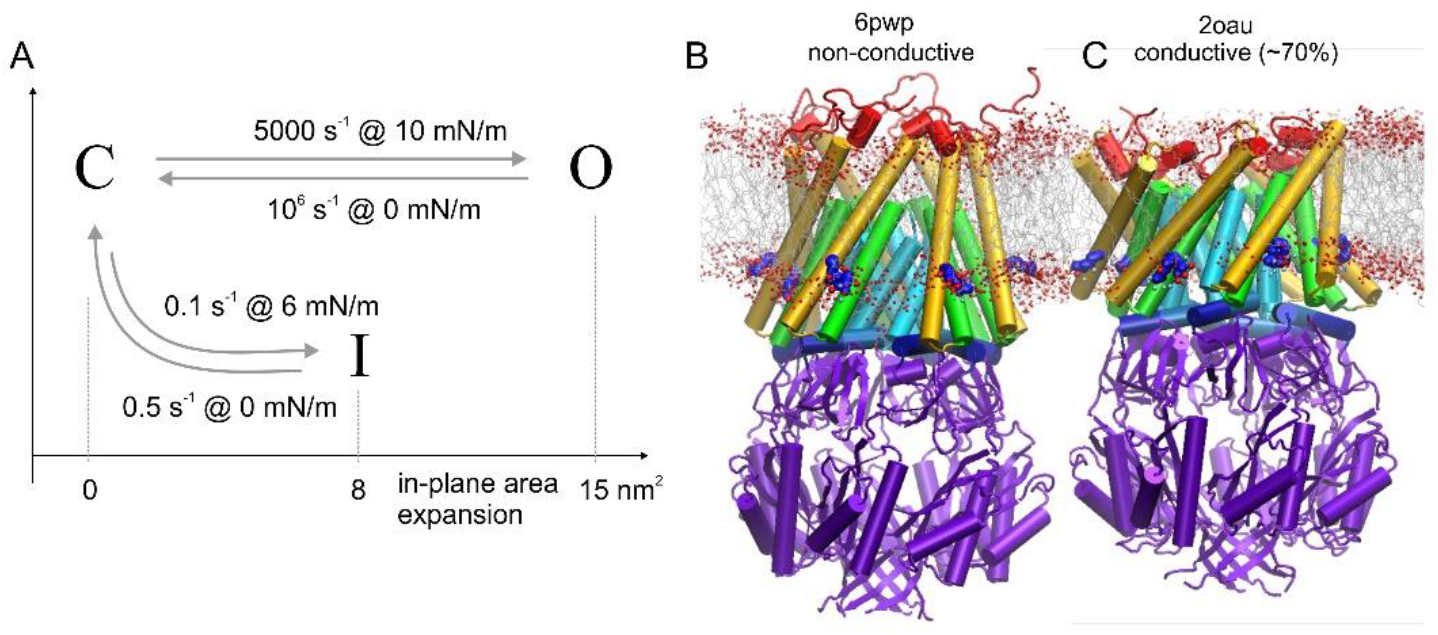
Description of the system. (A) The kinetic scheme of WT MscS with two pathways for the opening and inactivation transitions and the associated spatial and temporal scales. The opening and closing transitions (C↔O) are much faster and associated with a larger in-plane area change. 6PWP (B), the most complete non-conductive structure and 2VV5 (C), a conductive structure are both shown inserted in a simulated PE/PG bilayer (structures shown schematically) with the positions of R46 and R74 (blue sidechains) against the lipid boundary layer. Red dots represent lipid oxygens. Panels B and C represent the commonly referenced structural information on MscS which does not explain the kinetic scheme in panel (A).

Despite the fact that more than 40 crystal and cryo-EM structures of MscS have been deposited into the databases, it is still unclear how they relate to the open, closed, and inactivated functional states revealed in patch-clamp experiments (Belyy et al., 2010a) and in-vivo osmolyte release studies (Moller, 2023). Most of the published structures fall into two categories: (1) nonconductive and characterized by kinked pore-lining TM3 helices and splayed lipid-facing TM1-TM2 helical pairs; (2) semi-open with slightly straightened TM3 helices still separated from the lipid-facing helices and satisfying only about 65-70% of the fully open channel conductance. The simulated structures from these two different classes are presented in Figure 1B and 1C. Whether the protein adopts the first or the second category of structure seems to depend on the method of solubilization. Variations in the solubilizing conditions for identical protein preparations have revealed transitions from the semi-open to the non-conductive conical state that depend solely on the presence of specific detergents or lipids in the solubilization mixture (Flegler et al., 2021).

Indeed, lipid arrangement and the state of the membrane, especially around the protein cytoplasmic periphery, appears to play a significant role in structure stabilization. Splayed structures are characterized by lipid-facing TM1-TM2 pairs of helices uncoupled from sharply kinked TM3s and by deep crevices that are filled with lipids (.). MD simulations showed that the lipid bilayer can be strongly distorted in the vicinity of the splayed protein; lipids around the conical protein complex wedge between the helices, assume horizontal orientation and generally form non-bilayer structures (Pliotas et al., 2015).

The two structures depicted in Figure 1, panels B and C alone do not explain the three-state kinetic scheme presented in panel A. Towards modeling the MscS functional transitions, we look to leverage spatiotemporal parameters and other information in order to interpret existing structures. With the knowledge that there are two separate expansion pathways, we take advantage of the fact that at least one appears to involve substantial rearrangement of the protein-lipid boundary. We predict that conserved ‘anchoring’ residues located at the inner leaflet and interact with membrane phosphates should influence or be involved in the lipid rearrangement at the protein-lipid boundary. We, therefore, introduce mutations to probe which transitions are most sensitive to these substitutions. We anticipate that separate effects of mutations of canonical lipid-facing sidechains on the rates of the opening-closing and inactivation-recovery processes should be able to indicate the character of these transitions.

In the present paper, we focus on the role of two conserved arginines (R46 and R74) that prominently anchor the TM1-TM2 helical pairs to the boundary of the inner phospholipid leaflet (Fig. 1B,C). We show that the presence of these two charged anchors (14 per protein complex) imparts substantial voltage dependence to the processes of MscS adaptation, inactivation, and recovery. Replacement of these arginine pairs with aromatic residues retains channel functionality but removes voltage dependence. Tryptophan substitutions for R46 and R74 have a negligible effect on the closing rates but noticeably alter the rates of inactivation and recovery.

The data strongly suggest that the slow inactivation-recovery processes involve more substantial rearrangements at the protein-lipid boundary than opening and closing. The data also point to the role of metabolically-generated hyperpolarizing membrane potential in the regulation of osmolyte release valves by helping them to return from the dormant inactivated state back to the ready-to-fire state.

## Methods

### Mutagenesis

E. coli MscS wild-type (WT) and MscS mutants were expressed in the knockout strains MJF465 *(ΔmscL ΔmscS ΔmscK*) or PB113 *(ΔmscS ΔmscK*) from the IPTG-inducible pB10d vector, a modified pB10b plasmid (Iscla et al., 2004). MscS mutants were generated using the Q5 Site-Directed Mutagenesis Kit (New England Biolabs).

### Electrophysiology

Giant spheroplasts for patch-clamp electrophysiology experiments were generated as described previously (Martinac, 1987). Channel population recordings were made on excised inside-out patches in symmetric buffer containing 400 mM sucrose, 200 mM KCl, 50 mM MgCl_2_, 5 mM CaCl_2_, and 5 mM HEPES at pH 7.4. Membrane patches were obtained using borosilicate glass pipettes (Drummond) and recorded on an Axopatch 200B amplifier (Molecular Devices). Negative programmed pressure stimuli were applied using a high-speed pressure clamp apparatus (HSPC-1; ALA Scientific) using the Clampex software (Molecular Devices). Pressure and voltage protocol programming and data acquisition were performed with the PClamp 10 suite (Axon Instruments).

The patch responses were first recorded at +30 mV (pipette) under symmetric 1-s linear pressure ramp protocols to determine the pressure midpoints and saturating tensions for channel populations (Figure 2). These triangle ramps generate dose-response curves that were used for pressure to tension conversion for downstream fitting and analysis. Specifically, pressure to tension conversions were done by heterologously expressing MscS WT and mutants in the PB113 strain, which is devoid of all major mechanosensitive channels except MscL. Assuming the radius of curvature of the membrane patch (r) does not change, the ratio of the tension and the pressure midpoints between expressed MscS and native MscL should be constant according to the law of Laplace (γ=pr/2). In this way, MscL can be used as an internal standard with a known midpoint tension γ_0.5_ = 14 mN/m and known midpoint ratio of 0.6, enabling the midpoint tensions (γ_0.5_) for MscS mutants to be calculated (Belyy, 2010b). Traces recorded in the PB113 strain for WT and all mutants are shown in Supplemental Figure S1. The midpoint positions are marked with vertical bars and the ratios are listed in Supplemental Table S1. Ramp responses were fitted using QuB (Milescu et al., 2006; Qin et al., 1996)

**Figure 2.**
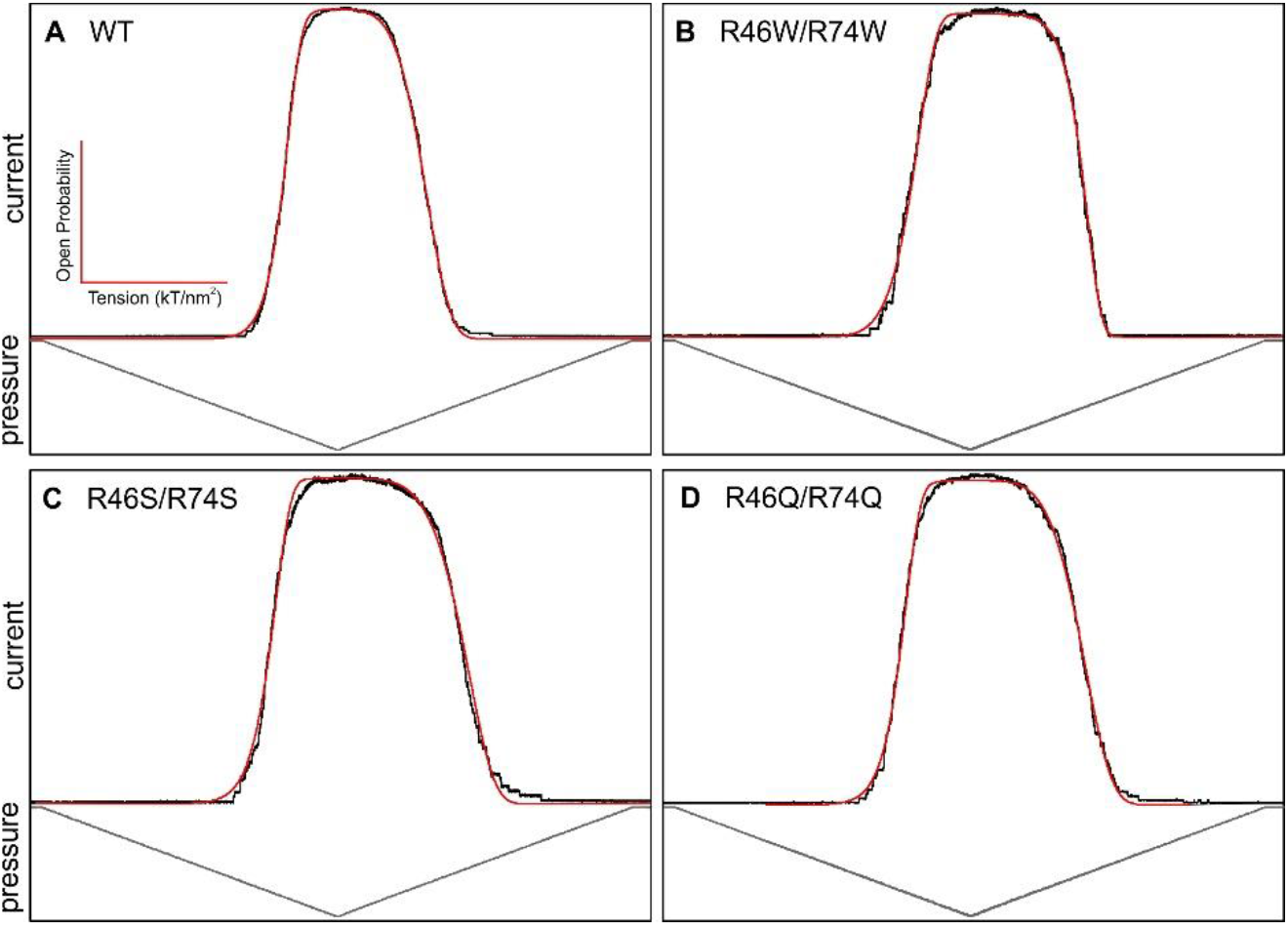
QuB fits of triangular pressure ramp responses. Channel population responses to symmetric 1-s triangular pressure ramps (grey) for (A) WT, (B) R46W/R74W, (C) R46S/R74S, and (D) R46Q/R74Q mutants (black). These dose-response curves were fitted with Qub (red) to obtain opening and closing kinetics (eqn 1). The ascending legs of each curve were also fitted to the Boltzmann distribution to obtain thermodynamic parameters (eqn 2). The extracted gating parameters are listed in Tables 1 and 2. (n=5)

Additional protocols were also used to extract kinetic and thermodynamic parameters associated with opening, closing, and inactivating transitions. Closing rate pulse-step protocols were performed to determine tension-dependent closing rate parameters (Figures 3, S3).

**Figure 3.**
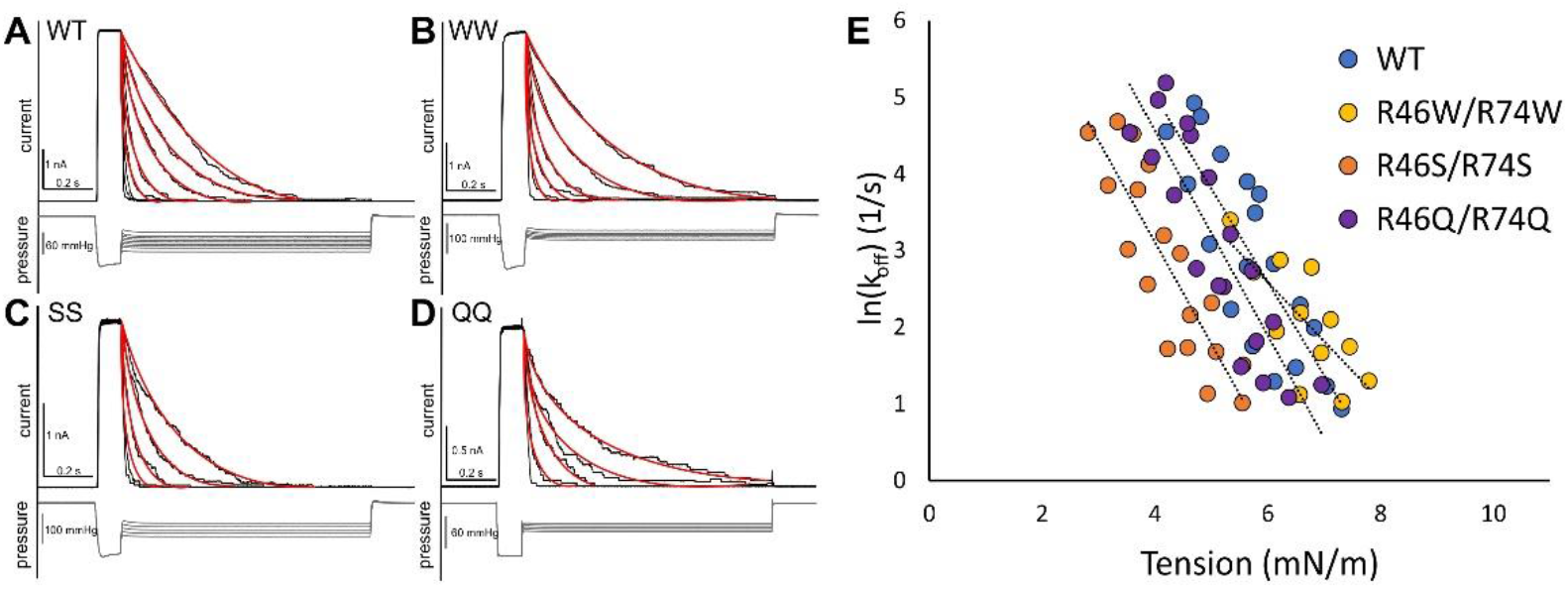
Effect of R46/R74 mutations on closing rate. The kinetics of the open → closed transition was probed using pulse-step protocols in patch clamp. (A-D) Channels were pulsed open and then allowed to close under various holding tensions (black). The closing behavior was then fit to the monoexponential function (red) for WT and all mutants. (E) A plot of ln[k_off_] vs tension yields a linear dependency where ΔA_OB_ and the closing rate at zero tension are given by the slope and y-intercept, respectively. Kinetic parameters are listed in Table 1.

Inactivation “comb” protocols in which channels closing under tension are periodically probed with saturating pulses to determine what portion of the population remains tension-sensitive (Akitake et al., 2005) were employed to monitor the rate of inactivation (Figure 4). Recovery from inactivation pulse-step-pulse protocols were also done by pulsing at saturating tensions to reveal the full population, holding at the midpoint pressure to encourage channels to close and inactivate, and then pulsing again with a series of saturating pressure pulses to extract the rates of recovery from the inactivated to the resting state (Figure 5). Minding the compromised stability of stressed patches, all experiments were performed across a range of voltages (±30,60 mV) to test the voltage-dependent character of these state transitions.

**Figure 4.**
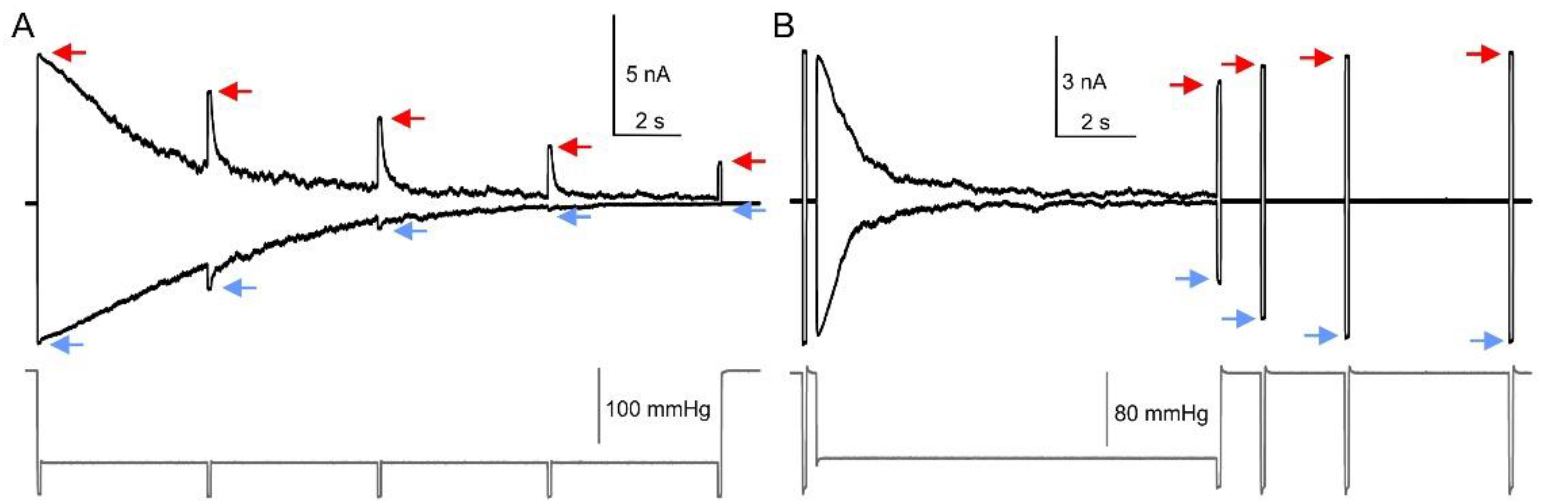
Patch-clamp protocols for measuring the kinetics of inactivation and recovery from inactivation. The protocols are illustrated for WT MscS. Patch clamp traces obtained using the “comb” protocol, which monitors the rate of channel inactivation by allowing channels to close under constant tension while intermittently pulsing with saturating pressure to check what portion of the population is still active and able to respond to tension. Fitting the decaying peak heights to the monoexponential function allows us to extract the kinetics of the inactivation process. (B) The kinetics of recovery from inactivation. The rate of recovery from the inactivated state back to the resting state can similarly be monitored by periodically pulsing the channels with saturating pressure to probe the population for activity following inactivation.

**Figure 5.**
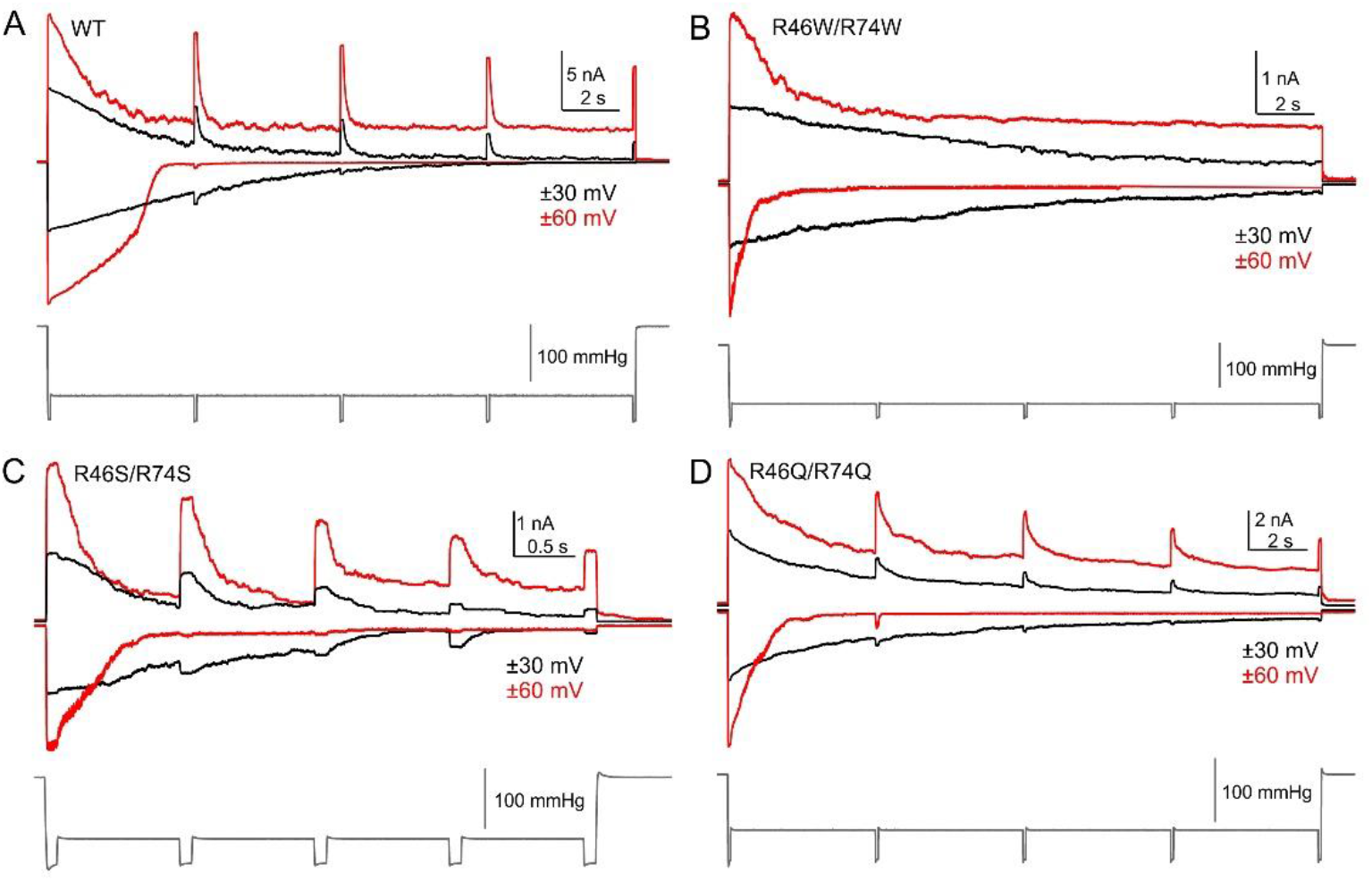
Rate of inactivation of MscS WT and mutants characterized in patch clamp at different voltages. The voltage displayed on the traces is the voltage held in the pipette, the effective cytoplasmic voltage is the opposite sign. The tips of the current responses to the saturating test pulses of the ‘comb’ pressure protocol reflect the kinetics of inactivation.

## Results

### Gating parameters from fitting traces to kinetic models

MscS channels residing in the *E. coli* inner membrane gate directly in response to lateral tension in the lipid bilayer (ref). This gating process can be studied in patch clamp, where patches of membrane containing channels are subject to pressure gradients that generate tension according to Laplace (Nomura et al., 2012; Sukharev et al., 1999). Thus, pressure stimulation on excised patches activates the channel population, producing a measurable current. When activated with saturating pressures, the full population of channels open (C→O). As applied pressure decreases, the population will gradually close in a tension-dependent manner (O→C). Under sustained moderate pressures, closed channels can transition into the inactivated state, in which they are nonconductive and tension-insensitive (C→I). It has been shown that WT MscS channels cannot inactivate from the open state, only from the closed state when subjected to moderate tension (Kamaraju et al., 2011).

The pressure protocols used in this study are capable of directly measuring the closing, inactivation, and recovery rates. The exception is the steeply tension-dependent opening process. It was shown previously that in the intermediate-to-high tension range, MscS opening can be faster than the speed of pressure clamping accessible with an HSPC-1 machine (Boer et al., 2011), and so resolving the tension-dependent kinetics of MscS opening directly may not be possible. For this reason, we retreat to recording population responses to 1-s triangle ramps, since saturating pressure applied over short (1-s) timescales precludes inactivation and results in a system that is essentially two-state (Figure 2). Population responses to triangular 1-s pressure ramps were fitted with QuB to a two-state kinetic model (O↔C) according to the Arrhenius-type relation:

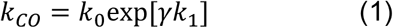

where k_CO_ is the transition rate, or the probability per unit time to make a transition from the closed to the open state, k_0_ is the intrinsic frequency with which the system overcomes the energy barrier between the closed and open states at zero tension; here γ is the applied tension, and k_1_ is the tension sensitivity. The closing rates can also be directly studied using the pulse-step protocol and exponential fitting (described below), which produces comparable results to QuB. We illustrate the pressure protocols and the quality of fitting procedures for WT MscS in Supplemental Figures S1 and S2, and S3. Main text figures show the comparison of responses between WT and mutants.

Thermodynamic parameters ΔA_CO_ and ΔE_CO_ were also determined from the Qub fitted parameters from the relationships: ΔA_CO_ (nm^2^) = (k_1_-k_1_’)(4.14 pN·nm); ΔE_CO_ (kT) = ln(k_0_/k_0_’). An analogous term for the tension sensitivity, ΔA_CO_, has a physical manifestation as the change in in-plane area of the protein in the membrane as it transitions between any two states, in this case the closed and open states. The Helmholtz free energy, ΔE_CO_, is the free energy of this transition in the absence of tension. Additionally, ΔA_CO_ and ΔE_CO_ were independently extracted from a Boltzmann fit of the opening leg of 1-s symmetric triangle pressure ramps for comparison. The Boltzmann distribution of channels under tension takes the form:

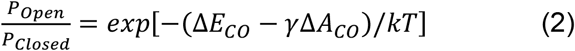

where ΔE_CO_ and ΔA_CO_ are as described, γ is the tension, k is the Boltzmann constant, and T is the temperature.

Similarly, the tension dependency (ΔA_OB_) of channel closing can be determined using a closing rate (pulse-step) protocol, in which the characteristic time of channel closure, specifically the tau τ, is extracted from a direct monoexponential fit of current decay as a function of tension (Figure 3A). A semi-log plot of tau versus tension yields a linear dependency, where ΔA_OB_ and intrinsic closing rate are the slope and y-intercept, respectively (Figure 3E). ΔA_OB_ represents the change in protein in-plane area between the open state energy well and the energy barrier between the open and closed state wells. Supplemental Figure S3 shows example traces and fitting for the closing rate protocol. Figure 3 shows characteristic traces and fitting for all mutants.

Compared to opening and closing (C↔O) transitions, inactivation and recovery transitions (C↔I) occur on much slower timescales, making them more tractable to determine experimentally. The rate of inactivation was determined via application of the ‘comb’ protocol, as described in the methods (Figure 4). While inactivation varies somewhat between batches, the process of recovery from inactivation, on the other hand, is easily observed with a series of short saturating test pulses that test for sensitive sensitivity following a conditioning step that drives the channel population to the inactivated state (Figure 5).

Regarding the mutagenized sites, we should explain that the conserved arginines 46 and 74 located on TM1 and TM2 are positioned at the same level facing the cytoplasmic phosphates. They apparently act as interfacial lipid anchors that stabilize the cytoplasmic ends of TM1 and TM2 helices in the membrane as the protein transitions between states. While single substitutions such as R46W or R74W produce similar but mild phenotypes, we decided to study double substitutions aiming at more discernable effects. A wide range of amino acid substitutions were made and tested that will be fully detailed in a coming paper. Here, we discuss the results of aromatic (W) and polar (S, Q) substitutions on inactivation and recovery as well as the effect of voltage on these transitions.

### R46/R74 mutations do not significantly affect the C↔O transition

The opening and closing phenotypes of WT and mutant channels were analyzed using patch clamp as described. Figure 2 shows characteristic traces of the full channel population in response to 1-s linear ramps of increasing pressure that drives channels to the open state, followed by decreasing pressure back to zero that allows them to close. Population saturation can be seen as a plateau at peak pressures. The result is a bell-shape that is slightly asymmetric due to closing hysteresis. The current was converted to open probability and the entire curve was fitted with QuB to yield kinetic parameters k_0_ and k_1_ for both C→O and O→C transitions (Table 1). Analogous fitting was done on the ascending leg of the triangle ramp according to Boltzmann’s distribution to extract thermodynamic parameters of the C→O transition, ΔA_CO_ and ΔE_CO_ (Table 2).

**Table 1.** Kinetic parameters for two-state fits. Kinetic parameters were obtained from fitting of 1 s triangular ramps with Qub (K_0_ and K_1_) and from the tension-dependent closing rate analyzed from patch clamp data (intrinsic closing rate), which are in good agreement with one another. The data show that closing behavior is fast, on the order of 10^5^, and that this fast closing is not significantly affected by mutations to R46/R74.(n≥3)

|  | C→O Transition |  | O→C Transition |  |  |
| --- | --- | --- | --- | --- | --- |
|  | K<sub>0</sub> (1/s) | K<sub>1</sub> | K<sub>0</sub> (1/s) | K<sub>1</sub> | Intrinsic Closing Rate (1/s) |
| **WT** | $(6.2 \pm 6.3) \times 10^{-7}$ | $2.2 \pm 0.1$ | $(6.3 \pm 4.9) \times 10^4$ | $-1.1 \pm 0.3$ | $(2.4 \pm 0.2) \times 10^5$ |
| **WW** | $(1.2 \pm 0.4) \times 10^{-5}$ | $1.2 \pm 0.0$ | $(2.4 \pm 3.9) \times 10^6$ | $-1.2 \pm 0.2$ | $(5.4 \pm 3.3) \times 10^5$ |
| **SS** | $(3.1 \pm 5.8) \times 10^{-6}$ | $2.0 \pm 0.4$ | $(3.1 \pm 5.5) \times 10^4$ | $-1.0 \pm 0.4$ | $(1.4 \pm 0.6) \times 10^4$ |
| **QQ** | $(3.0 \pm 2.4) \times 10^{-5}$ | $1.6 \pm 0.2$ | $(7.5 \pm 4.2) \times 10^3$ | $-1.0 \pm 0.2$ | $(1.2 \pm 1.2) \times 10^5$ |

**Table 2.** Spatial and thermodynamic parameters for two-state fits. Spatial parameters ΔA_CO_ describing the in-plane expansion of the channel upon opening, ΔA_OB_ describing tension sensitivity of closing, and ΔA_OB_/ ΔA_CO_ describing the position of the energy barrier between the closed and open states. [Need an energy diagram figure].(n≥3)

|  | C→O Transition |  |  |  | O→C Transition |  |  |
| --- | --- | --- | --- | --- | --- | --- | --- |
|  | Qub Fit |  | Boltzmann Fit |  |  | Qub | Boltzmann |
| | $\Delta A_{CO}$ (nm<sup>2</sup>) | $\Delta E$ (kT) | $\Delta A_{CO}$ (nm<sup>2</sup>) | $\Delta E$ (kT) | $\Delta A_{OB}$ (nm<sup>2</sup>) | $\Delta A_{OB}/ \Delta A_{CO}$ | $\Delta A_{OB}/ \Delta A_{CO}$ |
| **WT** | $14 \pm 2$ | $-25 \pm 2$ | $17 \pm 1$ | $-32 \pm 3$ | $7 \pm 1$ | $0.5 \pm 0.1$ | $0.4 \pm 0.1$ |
| **WW** | $10 \pm 1$ | $-25 \pm 2$ | $7 \pm 1$ | $-21 \pm 3$ | $6 \pm 1$ | $0.6 \pm 0.1$ | $0.9 \pm 0.2$ |
| **SS** | $13 \pm 2$ | $-24 \pm 4$ | $14 \pm 2$ | $-27 \pm 3$ | $6 \pm 1$ | $0.5 \pm 0.1$ | $0.4 \pm 0.1$ |
| **QQ** | $11 \pm 1$ | $-20 \pm 2$ | $11 \pm 1$ | $-23 \pm 3$ | $6 \pm 1$ | $0.5 \pm 0.1$ | $0.5 \pm 0.1$ |

An additional way to determine gating parameters for the O→C transition is to employ a closing rate (pulse-step) protocol consisting of a short pressure pulse to open channels followed by pressure steps to various sub-saturating levels as seen in Figure 3A-D. The kinetics of closure are determined as a function of tension and time, yielding the kinetic parameter of intrinsic closing rate at zero tension (Table 1) and the spatial parameter of ΔA_OB_, which gives information about the position of the energy barrier between the closed and open states (Table 2).

The kinetic data from both fit methods are in good agreement. The results show that MscS WT and all mutant channels are stably closed at zero tension, having intrinsic closing rate k_0,C→O_ values between 10^−7^ and 10^−5^. Spurious opening is extremely rare, which is consistent with the function of these channels as tight osmoregulators. The tension dependence of the rates, k_1,C→O_, was consistent between WT and polar mutants (SS, QQ), but slightly lower for the aromatic WW mutant, possibly related to its increased midpoint tension (Figure S1). As expected, once the gating stimulus of tension is removed, the closing rate is fast for all mutants. QuB parameters gave closing rate magnitudes between 10^3^ and 10^6^ s^-1^, while data from patch clamp closing rate exhibited much less variability, consistently resulting in closing rates on the order of 10^4^-10^5^ s^-1^. It was found that the backward transition (O→C) has approximately half the tension dependence as the forward reaction indicating a closer position of the open well to the transition barrier. (Table 1).

Thermodynamic terms for C↔O transitions were determined using both QuB and Boltzmann fitting regimes and compared. The expansion area, ΔA_CO_, calculated from both methods indicated again that SS mutant has a ΔA_CO_ value that is within error of WT, which has ΔA_OC_ of 14±2 and 17±1 nm^2^ for QuB and Boltzmann methods, respectively. On the other hand, the values for the WW and QQ mutants were noticeably smaller, exhibiting ΔA_OC_ between 7 and 11 nm^2^. Considering that the unitary conductances of WW and QQ are the same as WT, this suggests the possibility that they are pre-expanded but still closed. The change in area between the open state and the energy barrier ΔA_OB_ provides information about the relative location of the energy barrier between the open and closed states. The barrier position, ΔA_OB_/ ΔA_CO_, is relatively constant at the halfway point for Qub fitted parameters. The Boltzmann fitted parameters also give barrier position at approximately 0.5 for WT and all mutants except WW, which is heavily skewed by the small ΔA_OC_, resulting in a barrier position of 0.9, very close to the open well. This may be a sign of silent pre-expansion or artifact of non-homogeneity of the population described previously for MscL (Chiang 2004).

Energetically, there is some difference between QuB and Boltzmann methods. Analysis with QuB revealed ΔE values that were very similar for WT, WW, and SS, at ∼ -25 kT. The ΔE for QQ was somewhat lower, at only -20 kT. Conversely, Boltzmann analysis indicated ΔE values for all mutants ranging from WW at -21 kT to WT at -32 kT.

From the fitted parameters we constructed an energy diagram for a two-state tension sensitive channel with the in-plane area of the channel as the reaction coordinate (Figure XX). The data from both fit methods are comparable and overall show that substitutions at positions 46 and 74 do not hinder gating and do not significantly alter C*↔*O transitions.

### R46/R74 mutations do affect the C↔I transition

Under moderate tensions over prolonged time periods, MscS will inactivate and will not open in response to abruptly applied tension. The tendency to inactivate for WT, WW, SS, and QQ were probed with patch clamp pulse-step-pulse inactivation-recovery protocols. The difference between the magnitude of the current between the first pulse and the second pulse is the portion of the population that was inactivated. Although membrane potentials of -180 mV typically present in energized bacterial cells are inaccessible via patch clamp, even at -60 mV WT percent inactivation is already quite low at ∼ 30%. The percent inactivated steadily increases as the membrane potential depolarizes through voltages -60 mV, -30 mV, +30 mV, and +60 mV, indicating that the transition to the inactivated state is voltage-dependent. Under +60 mV, WT fully inactivates. We should remember that inactivation, like closing, is a tension-dependent process and that the extent of inactivation will vary accordingly. All pulse-step-pulse protocols were performed with the midpoint pressure (±10%) as the holding pressure for comparison.

Anchoring arginines at 46 and 74 appear to modulate voltage-dependent inactivation. The introduction of tryptophans leads to almost full inactivation that is independent of applied voltage; the percent inactivation of WW mutants at -60 mV is already 81%, compared to 30% for WT. Serine similarly exhibits increased inactivation at all voltages. Glutamine shows an intermediate effect. All mutants are eventually driven to 100% inactivation once +60 mV is reached (Figure 6A).

**Figure 6.**
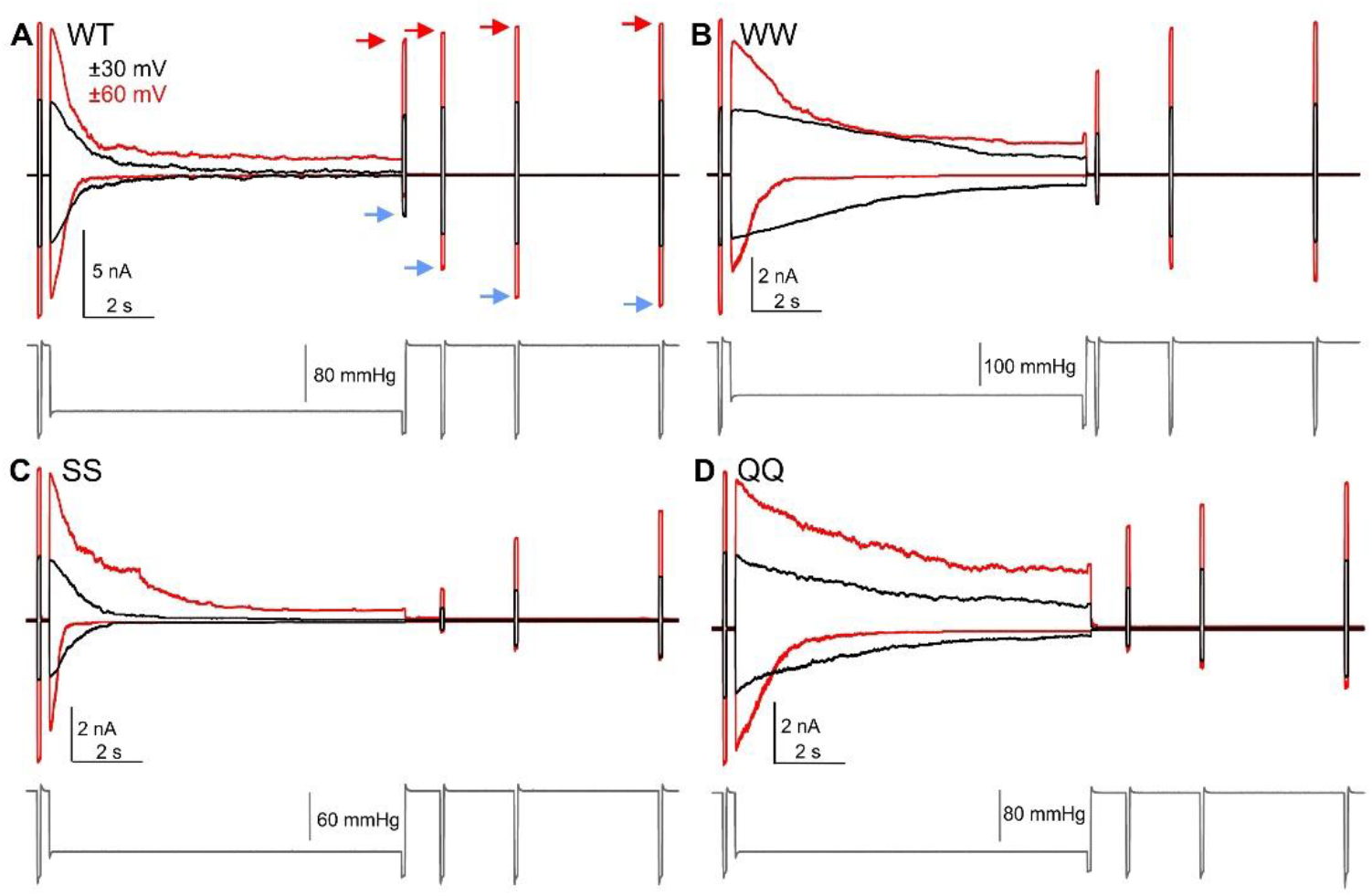
Voltage-dependent recovery from inactivation. The voltage displayed on the traces is the voltage held in the pipette, the effective cytoplasmic voltage is the opposite sign. (A) WT and (B-D) mutants were subjected to pulse-step-pulse inactivation recovery protocols to monitor the process of channel recovery from the tension-insensitive inactivated state to the tension-sensitive resting state. Channels are first pulsed open with saturating pressure to reveal the full population. Next, the channels open and adapt in response to lower conditioning pressures. After 10 s, channels are pulsed at saturating pressures again to determine the portion of the population that inactivated, followed by three more saturating pulses to observe the recovery process. For WT. both the extent of inactivation and the rate of recovery from inactivation are voltage dependent.

While C↔O transitions are fast, on the scale of ∼ 10^5^ s^-1^, transitions to and from the inactivated state are slower by 5-6 orders of magnitude. The rate of inactivation under midpoint tension, presenting here as the characteristic time tau, τ (s), varies from patch to patch, and the results for WT and mutants are statistically indistinguishable quantitatively (Supplemental Table 2B). Values for the characteristic time for all channels are on the order of 10^0^ to 10^1^ seconds.

However, the comb protocols used to observe the inactivation process reveal some qualitative characteristics (Figure 4). In WT, closing does not lead to immediate inactivation, as evidenced by the persistent peaks indicating that many channels remain sensitive to tension pulses over ∼ 20 s of applied tension at both -30 mV (top, black) and -60 mV (top, red) cytoplasmic voltages. Under positive voltages (bottom), WT channels close and immediately inactivate, showing no pulse activity from intermediate pressure pulses. Similar behavior was seen for SS and QQ mutants. WW mutants, on the other hand, displayed immediate inactivation upon closing even at negative cytoplasmic voltages.

Recovery back to the resting state from the inactivated state is likewise slow; the characteristic time (τ) is on the order of seconds. Since the inactivated state is immune to stress-activation, only the channels in the resting state can be activated by tension. Recovery from inactivation for WT has τ ≈ 1-2 s for all voltages. WW exhibits full inactivation and fast recovery that is significantly faster than WT, with τ ≈ 0.5-1s for all voltages. Conversely, SS and QQ substitutions slow recovery under both positive and negative voltages considerably, with τ ≈ 3-4 s and τ ≈ 2-10 s, respectively.

Figure 7 summarizes the inactivation kinetics parameters for all four versions of MscS studied here. The most drastic difference is observed between the WT channels showing the strongest voltage dependence and the WW mutant, the most voltage-independent. More specifically, at hyperpolarizing voltages (−60 mV, left side of the graphs), WT generally resists inactivation (5-25%) but gradually changes its behavior to full inactivation at depolarizing +60 mV. The WW mutant shows only a small variation between 80 and 100%, strongly favoring inactivation (panel A). The characteristic time of inactivation (tau) behaves non-monotonously, it is the longest for all mutants at hyperpolarizing -30 mV, but drops to 1 s for all mutants at + 60 mV. While WT shows the variation between 20 and 2 s, WW always ‘rushes’ into the inactivated state within 4 to a fraction of a second. Interestingly that WT and WW go hand-in-hand along the voltage scale in terms of tau of recovery, but WW always does it twice as fast. The small-sidechain double-serine mutant (SS) essentially retraces aromatic WW, whereas the large polar-sidechain QQ mutant shows intermediate behavior between WT and WW.

**Figure 7.**
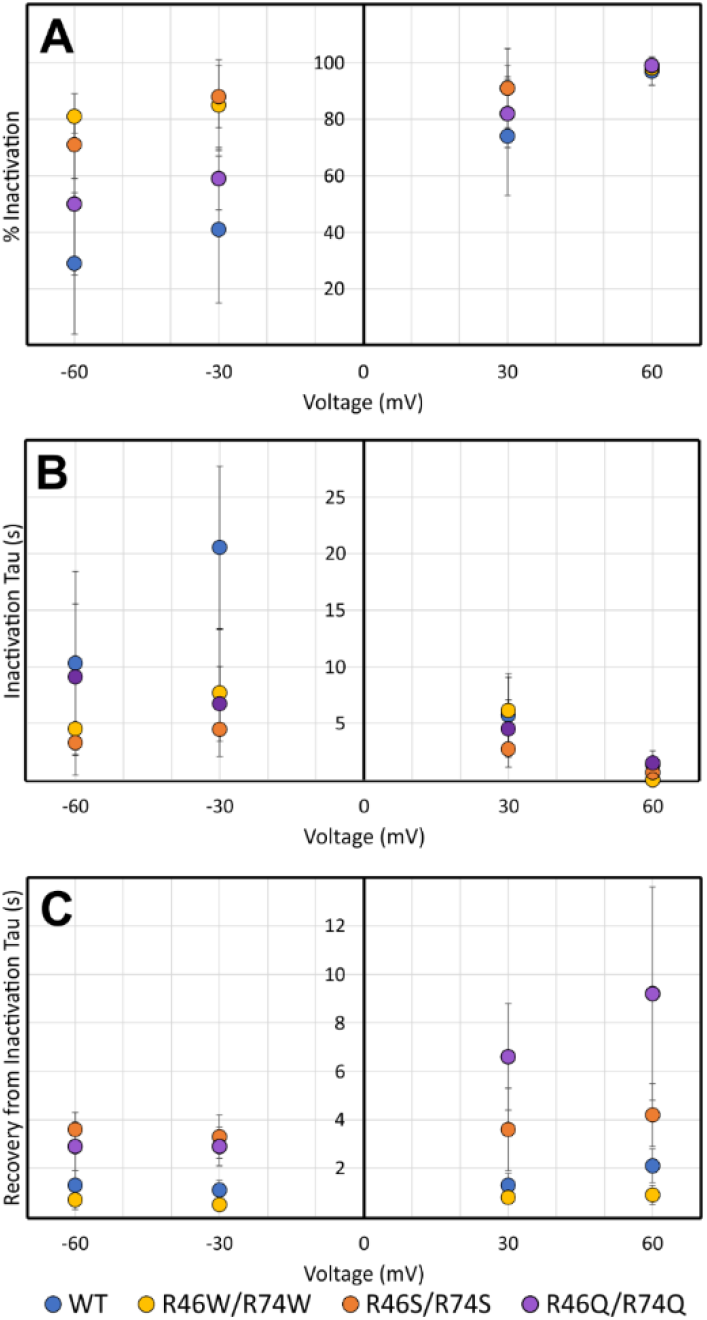
Summary of the voltage-dependent behavior of MscS WT and mutants. (A) The extent of inactivation. The percent of the channel population that inactivates under tension as determined from the pulse-step-pulse protocols shown in Figure 5 as a function of cytoplasmic voltage. WT is resistant to inactivation under hyperpolarized, physiological membrane conditions, but inactivates more readily as membrane potential depolarizes. WW and SS mutants almost fully inactivate independent of voltage. (B) The characteristic time, τ, of inactivation. WT slowly inactivates under negative voltage, but rapidly inactivates under positive voltage. WW and SS mutants inactivate quickly more or less irrespective of voltage. (C) The characteristic time, τ, of recovery from the inactivated state.

## Discussion

When the first crystal structure of MscS (2OAU) solved in foscholine (Bass 2002) was published, the abundance of arginines on the lipid-facing helices was noted and the TM barrels were positioned such that R46 and R74 were placed in the middle of the membrane. At that time, the complexity of the channel’s gating mechanism was not known and these arginines were predicted to impart voltage-sensitivity to the opening transitions (.). Later, the opening/closing transitions were shown to be almost voltage-independent and the presence of a prominent inactivated state was demonstrated (Akitake).

Results of previous simulations of MscS models (Anishkin) and more resent cryo-EM structures of MscS in nanodiscs (6PWP, 6PWN, 6UZH, 6YVL, 6YVM, 6YVK, 8DDJ) (Angiulli et al., 2020; Park et al., 2023; Reddy et al., 2019; Zhang et al., 2021) revealed that R46 and R74 are not in the middle but rather face the cytoplasmic boundary of the membrane interacting with lipid phosphates. This new placement of the channel relative to membrane boundaries strongly suggested that these arginines fulfill the anchoring function for TM helices during transitions and likely facilitate the process of protein folding and assembly according to the “positive inside” rule (refs).

From the experimental data presented above, we may conclude the following. The steeply tension-dependent C↔O transitions are 4-5 orders of magnitude faster than the C↔I transitions; correspondingly, at zero tension the closing rate is 6 orders of magnitude faster than the recovery rate.

The R46W/R74W substitutions do not substantially change the intrinsic closing rate, barely two-fold (Table 1), but do significantly change the propensity toward inactivation as well as inactivation and recovery kinetics (Fig. 7). The R to W substitutions maintain channel functionality, but essentially eliminate the voltage dependence of both inactivation and recovery. Effects of polar uncharged residues in these positions depend of the size of the sidechain and its H-bonding capacity. The small sidechain serine slows down the rate of recovery and makes it completely voltage-independent. The large-sidechain glutamine slows down the recovery even more, but retains some voltage-dependence.

The presented phenomenology begs for a mechanistic explanation. The three key questions posed by the described phenomenology that need to be addressed are: (1) why is the opening transition essentially voltage-independent? (2) how does voltage modulate the stability of the inactivated state and why are the charge and polarity of sidechains in positions 46 and 74 necessary for this modulation? (3) what is the role of interfacial anchors in assisting the tension-driven transitions?

Answers to all three questions will require extensive Molecular Dynamics simulations of existing structures and models under different tensions and voltages. The R46W/R74W mutant needs to be carefully compared to WT MscS in terms of stability and the structure of the protein-lipid boundary as well as in terms of the distribution of electrostatic potential in the entire system under different voltages. The last question on how the lipid ‘anchors’ work will employ the concept of ‘solvation by lipids’ and will involve extensive analysis of favorable contacts of canonical anchoring sidechains with lipids in different conformations. This will be a modification of the allosteric model of tension-driven transitions in membrane proteins accounting for differential exposure of sidechains with different affinity to lipids.

And finally, from the biological point of view, voltage-dependence of inactivation and recovery points to a potentially novel mechanism of regulation of MscS-type osmolyte release valves by natural metabolically-generated membrane potential that exists and works as an energy-coupling intermediate in essentially all bacteria utilizing the electron-transport chain. In fully energized *E. coli*, membrane potential is estimated to be near -180 mV (refs). This potential should keep most of the MscS population in the closed ‘ready-to-fire’ state. If a bacterium is starved and cannot maintain membrane potential, part of the population may go to the dormant inactivated state to exclude spurious openings and loss of precious metabolites. Through the membrane potential, the cell will exert its control over the metabolite release valves making them dependent on the metabolic state of the cell.

## Acknowledgement

This work was supported by the NIH R01AI135015 to S. Sukharev and by the National Science Foundation Graduate Research Fellowship Program under grant no. DGE 1840340 to E. Moller. Any opinions, findings, and conclusions or recommendations expressed in this material are those of the author(s) and do not necessarily reflect the views of the National Science Foundation.

